# CRISPR screening reveals SYCP3 as a key driver of metastasis in prostate cancer

**DOI:** 10.1101/2025.01.30.629925

**Authors:** Maria Rodrigo-Faus, Ines del Monte-Garcia, Marina Hermosilla-Trespaderne, Alicia Gordo-Vega, Natalia Vidal, Javier Puente, Melchor Saiz-Pardo, Ángel M Cuesta, Hui-Qi Qu, Hakon Hakonarson, Almudena Porras, Daniel Sanchez-Parcerisa, Paloma Bragado, Álvaro Gutierrez-Uzquiza

## Abstract

Prostate cancer (PCa) is a prevalent male cancer with high survival rates, except in advanced or metastatic stages, for which effective treatments are lacking. Metastatic PCa involves complex mechanisms including loss of tumor suppressor genes and DNA repair molecules, which impacts therapy responses. We have reanalyzed data from a CRISPR/Cas9 genome wide screening previously performed to identify essential regulators of invasive abilities of the metastatic cell line DU145 identifying *SYCP3* as a regulator of metastatic invasion. Subsequent analyses of tumor samples demonstrated that *SYCP3* expression is frequently upregulated in PCa tumors from patients in advanced stages. Furthermore, *SYCP3* genetic depletion significantly reduced the invasive and migratory abilities of DU145 cells and increased their adhesion capacity. Additionally, and due to the implication of *SYCP3* on DNA repair processes, we have analyzed the role of *SYCP3* on the cellular response to radiotherapy (RT) and found that its depletion induced RT resistance, suggesting a role for *SYCP3* in DNA damage response and genomic instability. All these data support a role for *SYCP3* in PCa metastasis and provides opportunities for personalized medicine.

## INTRODUCTION

Prostate cancer (PCa) remains the most common solid tumor in men and the third leading cause of cancer deaths among men in the European Union, and its incidence increases rapidly with age ^1^. Although the high majority of PCa patients have a high 5-year survival rate ^2^, the survival rate of patients with an advanced or metastatic stage of the disease is severely decreased. Despite the several treatment options available for localized PCa tumors, no curative treatment has been developed yet for metastatic prostate cancer (mPCa). mPCa tumors show a high rate of gene loss, particularly tumor suppressor genes and molecules involved in DNA repair processes, which could be relevant to predict the outcomes to therapies^3–5^. PCa arises as a hormonally driven disease that depends on androgen receptor (AR) activation. It often progresses to become resistant to androgen deprivation therapy, leading to Castration-Resistant Prostate Cancer (CRPC), which can further evolve to metastatic CRPC (mCRPC)^5^ . There is still a small understanding of the molecular mechanisms that regulate the growth and progression of these mCRPC tumors considered highly treatable although unfortunately incurable^3,5^; in contrast to PCa primary tumors that can be optimally managed.

Metastasis is the main cause of death in people with cancer. To colonize distant organs, circulating tumor cells must overcome many obstacles through mechanisms that we are now starting to understand ^6^. Several key metastatic genes have been identified through candidate-gene approaches, such as genes coding extracellular–matrix-degrading metalloproteinases (MMPs), cathepsins or proteins involved in cytoskeletal rearrangements^7–10^ . However, targeted therapies directed against these genes have not shown to be effective at preventing the dissemination of cancer cells, supporting the fact that metastatic cells are able to overcome the blockage of these pathways through additional genes that remain to be discovered.

Since the survival of patients with metastatic prostate cancer is so low (approx. 3 years) ^11^, it is extremely important to establish whether newly emerging essential genes or factors controlling metastasis might offer new opportunities for personalized therapy. Genomic analysis and preclinical studies have found frequent genomic alterations in genes like *AR, TP53, PTEN, BRCA2/1* and epigenetic regulators^3^ in mPCa. These studies have also uncovered prognostic biomarkers and new mechanisms involved in therapy resistance ^12^. Approximately, 80% of the tumors contain co-occurring mutations, which include genes encoding transcription factors or regulators of the transcription^3^ . Additionally, 20% of mCRPC have somatic alterations in genes encoding molecules involved in DNA repair, such as *BRCA2* ^3–5,13^ . There is also an extensive list of less frequent alterations in mCRPC, whose predictive or prognostic value role remain poorly understood. The identification and study of these co-occurring alterations and less frequent alterations could provide a valuable insight for improving clinical risk stratification and developing new therapeutic strategies.

To identify effective therapeutic targets or biomarkers for mPCa, it is essential to examine the highest possible number of genes simultaneously, while minimizing bias. Genome wide studies have proven to be an effective strategy for this purpose. By using genome-wide libraries in the proper cellular models, researchers have been able to identify new therapeutic targets and/or biomarkers with prognostic value ^14–16^. Our research reanalyzed CRISPR/Cas9 data targeting ∼20,000 genes screening previously performed in our lab ^16^ and identified *SYCP3* as an essential gene for mCRPC cell invasion in the highly invasive, androgen independent PCa cell line DU145. Furthermore, our studies have found that *SYCP3*, a component of the synaptonemal complex involved in segregation of chromosomes in meiosis and strongly associated with DNA repair mechanism ^17^, is also a key factor for mPCa cell invasion, adhesion and resistance to radiotherapy.

## RESULTS

### *In vitro* CRISPR/Cas9 screening reveals several genes and biological processes significant for mPCa invasive processes

Since the progression of PCa disease to a metastatic stage is a complex process, we decided to identify new essential genes in mCRPC reanalyzing an unbiased high-throughput CRISPR/Cas9 screening targeting 20,000 genes (GeCKO V2 CRISPR knockout pooled library) already performed in two independent mPCa cell lines ^16^. Of the 2 CRISPR screenings performed and reported, we focused on the results obtained from the DU145 mPCa cell line with brain tropism, as brain mPCa has the lowest survival rate. Rodrigo-Faus and colleagues identified 884 genes that support the metastatic abilities (pValue<0.05 and gRNA>1) of mCRPC DU145 cell line and significantly reduced the invasive capacities of DU145 cells according to the MAGECK analysis^16^ (Fig 1A and Supplementary Table 1). The top 10 genes with higher MAGeCK scores are shown in Table of Fig. 1B. Notably, among the 884 genes, we found genes such as *WNT5A* ^18^ and *GP2* ^19^ that were previously associated to mPCa in previous studies, proving the validity of CRISPR/Cas9 libraries technology to unbiasedly identify relevant mPCa modulators.

**Figure 1.**
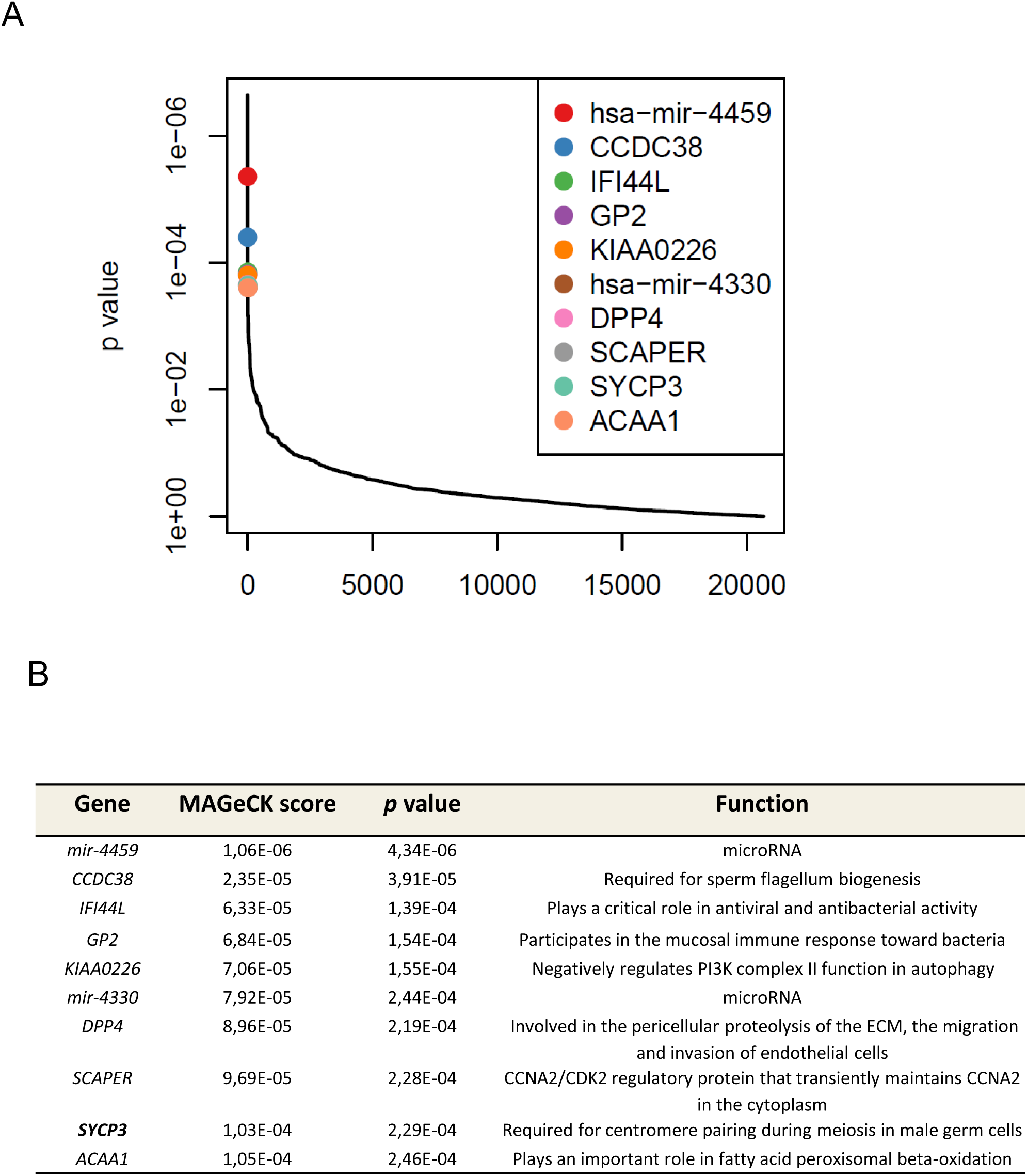
CRISPR/Cas9 screening reveals several key genes in the invasion process of PCa metastatic cell line DU145. **A** The positively selected genes associated with invasive process (y-axis: p-value; x-axis: gene). **B** Top genes significantly associated with mPCa invasive process in DU145 cells including their MAGeCK score and p-value in the screening and function.

We performed GO enrichment analyses of significant genes (Fig. 2) and observed several biological pathways enriched in pathways that impair metastasis such as nucleotide excision repair (Fig. 2A) or promote metastasis such as regulation of the MAPK cascade or G2/M transition of mitotic cell cycle (Fig. 2B). Moreover, gene set enrichment analyses (GSEA) showed enrichment in processes that either impair metastases such as hedgehog or mTOR pathways (Fig. 2C) or promote metastasis such as WNT/β-catenin signaling (Fig. 2D). Interestingly, several of these gene functions identified are associated with cancer progression and metastasis ^20^.

**Figure 2.**
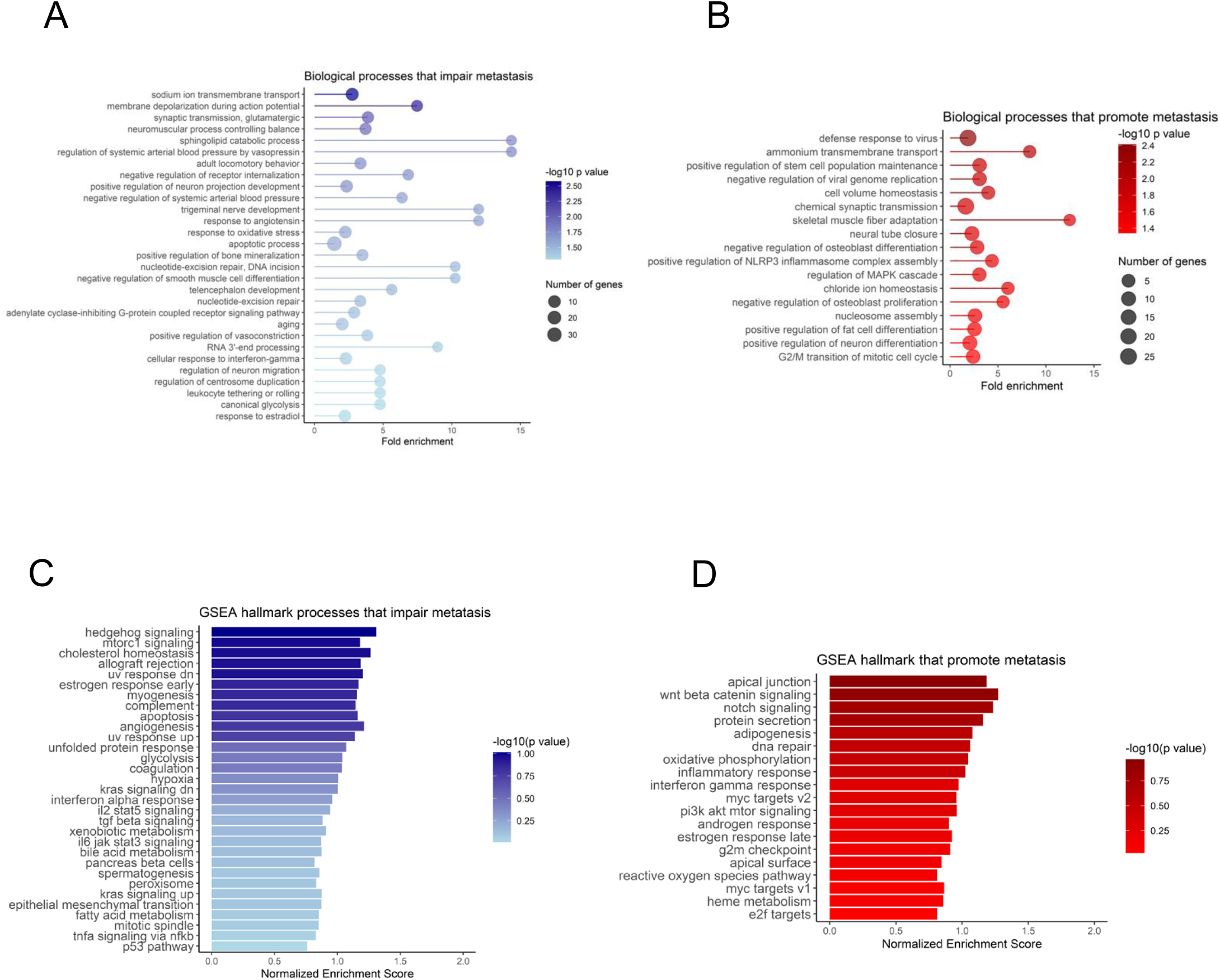
Gene ontology analysis reveals biological pathways important for PCa metastasis to the brain. **A-B** GO biological pathway enrichment analysis (GO:BP) enrichment for DU145 screening results. Biological processes that impair metastasis (A) or promote (B) metastasis are shown. **C-D** GSEA hallmark processes that impair (C) or promote (D) metastasis.

Remarkably, one of the top identified candidate genes was *SYCP3* that codifies synaptonemal complex protein 3 (SYCP3) which has been previously associated with DNA damage ^17^. *SYCP3* displays a specific expression restricted to spermatic-meiotic cells (Supplementary Figure 1) during the meiotic process to regulate the synaptonemal complex^21^. It is associated to chromosome cores at diplotene phase and to the sister centromeres during metaphase I ^22^. SYCP3 is a centromere-associated protein that dissociates from the centromeres at anaphase II and is not found in mitotic metaphase centromeres, conferring *SYCP3* a highly controlled expression in timing and cell type ^23^. Despite its restricted expression in physiological conditions, ectopic expression of *SYCP3* has been reported in several tumors unrelated to its natural expression ^24^.

### SYCP3 is upregulated in metastatic prostate cancer patients

To shed some light on the role of *SYCP3* in metastasis and dissemination we decided to investigate its expression in PCa tumors. First, we analyzed publicly available cancer databases to compare *SYCP3* mRNA expression levels between normal (N), primary tumor (PT) and metastatic (M) samples. We observed that *SYCP3* levels were upregulated in metastatic compared to primary tumor samples in three independent expression analysis datasets (Grasso, Taylor and Varambally) retrieved from CANCERTOOL database^25^ (Fig. 3A). Additionally, we also explored the role of *SYCP3* in PCa progression by analyzing the correlation between *SYCP3* expression and Gleason Score (GS). GS ranks the malignancy of PCa tumor cells according to the two main architectural patterns of tumor tissues ^26^. Analysis of *SYCP3* expression showed higher expression in tumor samples with high GS (>8) in two independent databases (Tailor and TCGA; retrieved from CANCERTOOL and UALCAN^25,27^ database respectively) (Fig. 3B and 3C, respectively). To confirm these results, we decided to analyze a cohort of FFPE-embedded PCa primary tumor samples recruited from Hospital Clínico San Carlos (Madrid, Spain). Tumor samples were divided into two groups according to their aggressiveness and the diagnosis of metastasis in the patient. In the group labeled as “GS 6”, tumor samples with a GS of 6 and no metastasis present were included, meanwhile samples with a GS equal or greater than 7 and metastatic at diagnosis were included in the group labeled as “GS≥7”. As Figure 3D shows, patients with higher GS displayed higher expression of *SYCP3* than patients with less aggressive tumors. Supplementary Figure 2 summarizes the characteristics of the patients included in the study.

**Figure 3.**
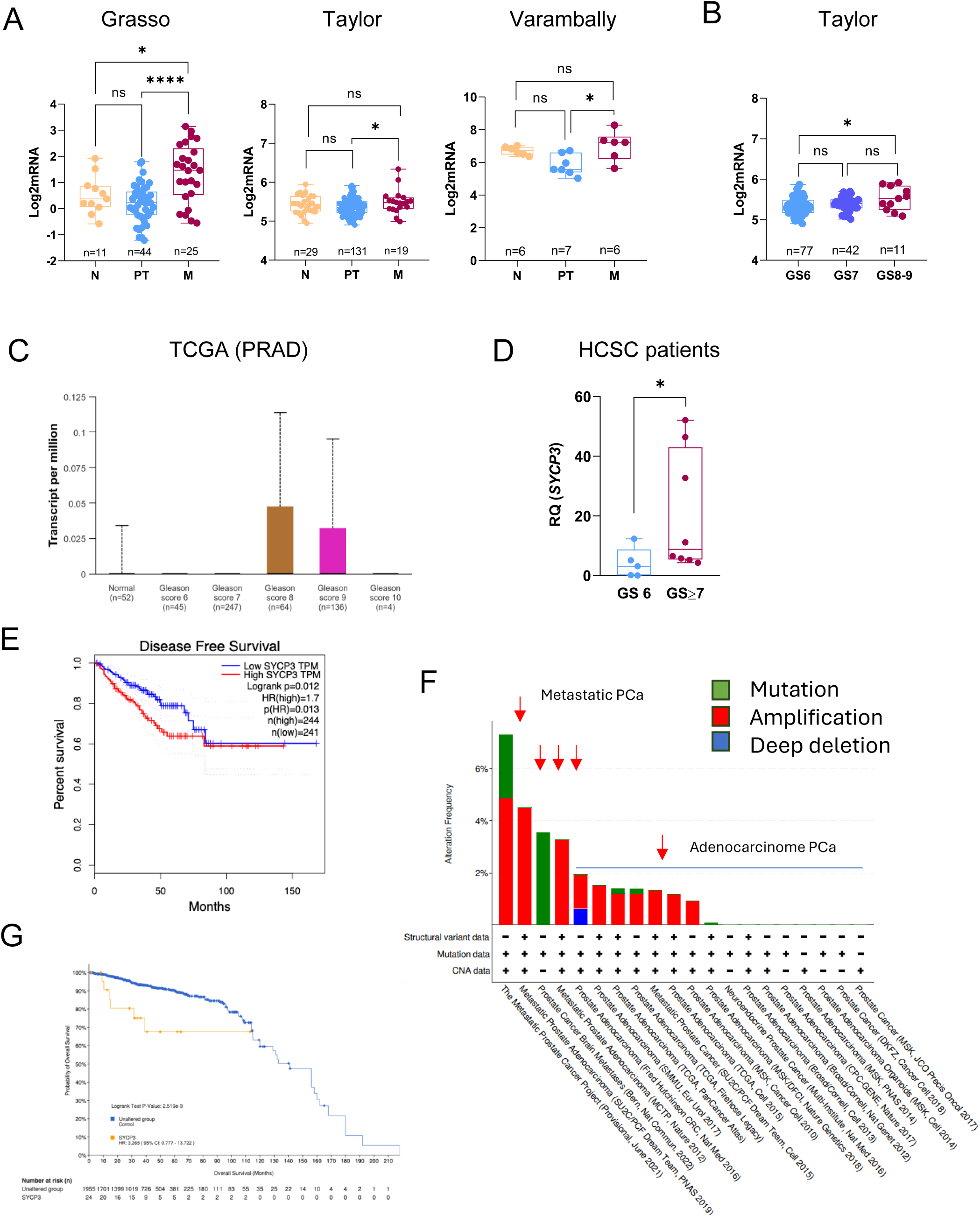
*SYCP3* is upregulated in PCa patient samples and predicts a poor diagnosis. **A** *SYCP3* mRNA levels in mPCa (M), primary tumor PCa patients (PT) and normal tissue (N) based on sample type obtained from Grasso, Taylor and Varambally cohorts of PCa patients obtained from CANCERTOOL database. Number of patients included in each group are depicted in the graph. **B-C** *SYCP3* mRNA levels based on GS in PCa patients from Taylor cohort obtained from CANCERTOOL database. Number of patients included in each group are depicted in the graph. **D** *SYCP3* mRNA levels in PCa patients’ samples obtained from Hospital Clínico San Carlos cohort based on GS (GS≥7 (GS≥7, grey color) or GS 6 (GS=6); n=13). **E** Kaplan–Meier curves showing the difference in disease free survival (DFS) between patients with high (red line) and low (blue line) expression levels of *SYCP3*. **F** Genetic Alteration frequencies with mutation type for *SYCP3* in PCa patient samples reported on indicated studies and obtained from cBioportal database. X-axis represents the type of alteration (red—amplification; blue— deep deletion; green—mutation) and the Y-axis represents the frequency of the alteration in different cancers. **G** Correlation between mutation status and overall survival of cancer patients in SYCP3. The orange line shows the overall survival estimates for patients with an alteration in the gene as compared to patients with no alteration (blue line). Statistic tests: One-way ANOVA test (**A-C**) U-Mann Whitney test (**D**) and Log-rank test (**E** and **G**). *, *P* < 0.05; **, *P* < 0.01; ***, *P* < 0.001; ****, *P* < 0.0001.

To investigate if *SYCP3* expression correlates with poorer outcomes, we analyzed the association between disease free survival (DFS) of patients and *SYCP3* expression using TCGA data. Kaplan–Meier method was used to compare DFS of patients with *SYCP3* expression, as described in the materials section. *SYCP3* mRNA expression across all PCa tissues was used to classify all cases into high (n = 244) and low *SYCP3* expression (n = 241). Figure 3E shows that patients with higher expression of *SYCP3* present shorter DFS compared with those with lower *SYCP3* expression (HR 1.7, *p* = 0.013).

All these results suggest that *SYCP3* expression is upregulated in PCa tumor samples and correlates with poor prognosis. These results, in combination with *SYCP3* restricted physiological expression, makes this gene a potential biomarker of PCa metastasis.

### SYCP3 is frequently altered in patient samples

To understand the molecular mechanisms behind the *SYCP3* upregulation observed in samples from PCa patients, we studied the prevalence of their genetic alterations across various human prostate cancers datasets, using the “Prostate adenocarcinoma” set of cBioPortal database (ICGC/TCGA, Nature 2020, available at http://www.cbioportal.org, accessed on July 5^th^, 2024). As shown in Figure 3F, *SYCP3* is altered in several prostate cancer studies. Interestingly, the highest alteration frequency observed in *SYCP3* (up to 6%) was present in samples from metastatic prostate cancer patients, being amplification the primary alteration type. A pan-cancer analysis on the same database showed that *SYCP3* also underwent amplification in various sets of tumor tissues (B-cell cancer, lung cancer, pancreatic cancer, cancer renal cell patients) (Supplementary Figure 3). Interestingly, although “amplification” was the most frequent alteration, “mutation” also appeared as the unique form of alteration in patients with brain-metastasized PCa (Fig. 3F), suggesting that missense mutations may contribute to this phenotype.

To identify potential genetic hotspots, we analyzed the mutation patterns affecting the *SYCP3* locus. As shown in Supplementary Figure 4, mutations in *SYCP3* gene appear to be randomly distributed across the entire sequence, without clustering in any critical region for protein activity. Finally, to confirm that genetic alterations in *SYCP3* impact PCa progression, we analyzed the association between *SYCP3* alterations and overall survival (OS) in PCa patients using the cBioportal dataset. The survival ratio of the “unaltered group” (without genetic alterations on *SYCP3* gene) was higher than the “altered group” survival (HR 3.27 (95% CI 0.8-13.7); p value 0.0025), which implies that *SYCP3* genetic alterations (mainly amplification) have an impact on the poor overall survival and prognosis in PCa (Fig. 3G).

### SYCP3 depletion decreases tumor cell motility and adhesion

*SYCP3* expression has already been associated with several cancers including acute lymphoblastic leukemia, ovarian tumor, brain tumor cancer and non-small cell lung cancer metastasis^24,28^. To assess whether *SYCP3* upregulation was also involved in the invasive properties of PCa cells as suggested by our CRISPR/Cas9 screening (Fig. 1), we decided to deplete *SYCP3* in the highly metastatic DU145 cell line using CRISPR approaches. A genetic perturbation was introduced in *SYCP3* locus altering the open reading frame (ORF) (Fig. 4A). Genetic deletion using Inference of CRISPR Edits (ICE) analysis showed that *SYCP3* locus was efficiently altered generating indels with high KO% in the ORF, while no effect on *SYCP3* was detected in non-targeting control (NTC) (Fig. 4A and Supplementary Figure 5). Using this cellular model, we evaluated the effect of *SYCP3* depletion on invasion using Matrigel-coated Boyden chambers using 10% FBS as chemoattractant. Our analysis found that invasion was reduced in SYCP3-KO cells as compared to NTC control cells (Fig. 4B) validating our initial screening results. This suggests that *SYCP3* expression promotes mPCa cell invasiveness. To confirm that cells showed and impaired migratory/invasive phenotype we performed a wound healing assay (Fig. 4C). In agreement with results from the invasion assay, the motility and ability to close the wound of *SYCP3* depleted cells was reduced compared to NTC control cells. Hence, these results indicate that *SYCP3* gene depletion produces a significant reduction in DU145 cell motility.

**Figure 4.**
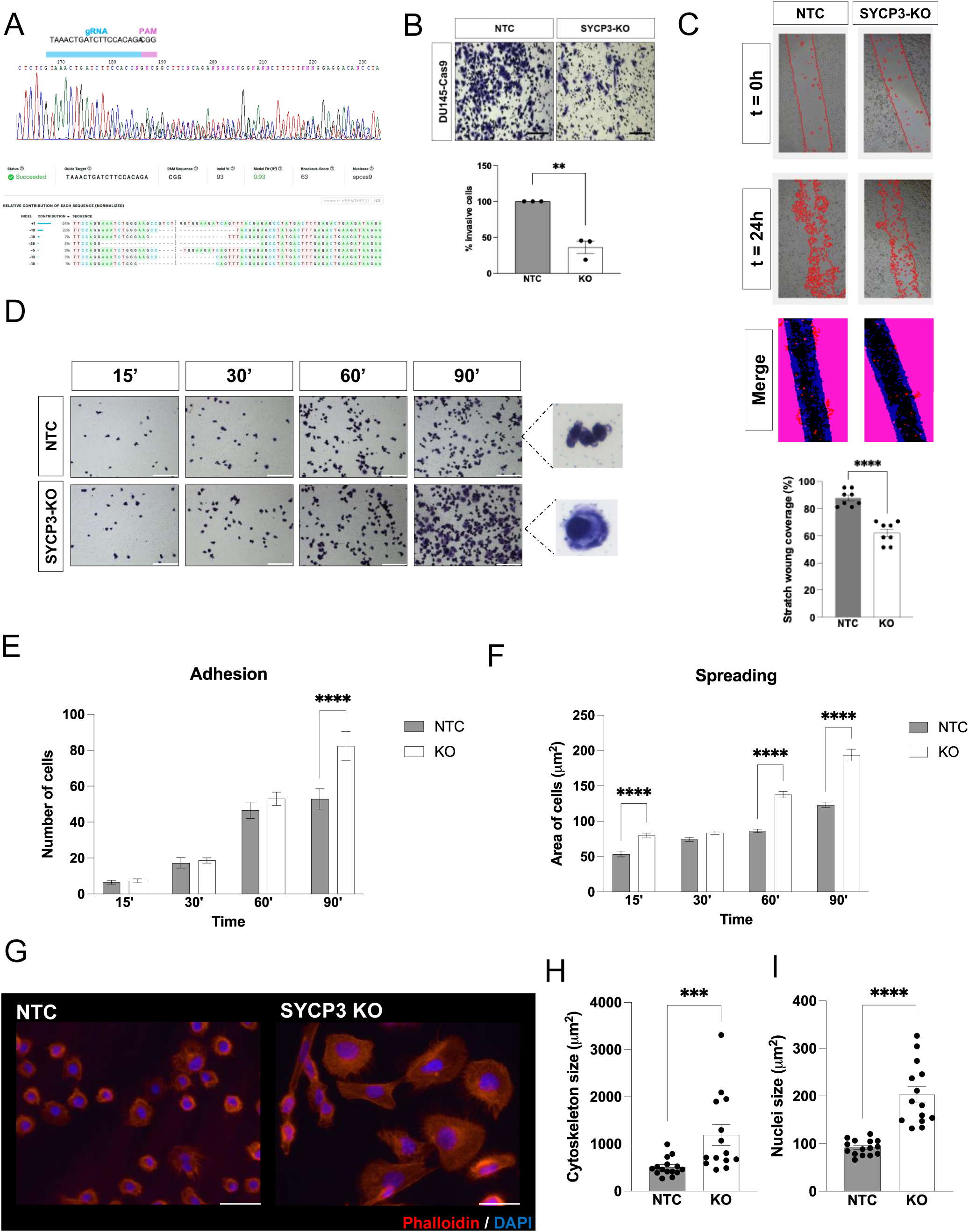
*SYCP3* genomic depletion reduces invasive abilities while enhances adhesion in PCa cells DU145. **A** The Inference of CRISPR Edits (ICE) software output of the analyses of the Sanger sequencing data on the *SYCP3* gene part flanking the exon 1 of the ORF. **B** Invasion analysis in DU145 cells using 10% FBS as a chemoattractant. Top panels, representative images of cells (bars: 100 µm); bottom panels, histograms showing the mean value ± S.E.M. of the percentage of invasive cells (n = 3). **C** Migration analysis of DU145 cells using a wound healing assay (n=3). **D-F** Adhesion/spreading analysis of DU145 cells. Representative images of adhered DU145 cells (C) and histograms showing the mean value ± S.E.M. of the percentage of adhered cells (D) or areas covered (E) of SYCP3 depleted cells referred to NTC cells (n = 3). Scale bars: 100 μm. **G** Immuno-fluorescence microscopy images of phalloidin staining (*red*) adhered NTC and SYCP3 depleted DU145 cells. Cell nuclei were stained with DAPI (*blue*). Scale bars: 100 μm. **H-I** Histograms showing the cytoskeleton (G) or nuclei area value ± S.E.M. of adhered DU145 indicated cells (n=2). Statistic tests: Student t test (**B, C**), two-way ANOVA (**E, F**) and U-Mann Whitney test (**H, I)**. *, *P* < 0.05; **, *P* < 0.01; ***, *P* < 0.001; ****, *P* < 0.0001.

We have demonstrated that *SYCP3* plays a role in promoting cell motility. To get further insight into the functions of *SYCP3* on mPCa, we analyzed the impact of *SYCP3* depletion on cell adhesion -a process highly connected with invasion. As shown in Figure 4D, the number of adhered cells increased in *SYCP3* depleted cells compared to NTC cells (Fig. 4D and E) after 90 minutes. Interestingly, we observed morphological differences on the shape of the cell between NTC and SYCP3-KO cell (Fig. 4F). Therefore, we evaluated the role played by *SYCP3* in cellular spreading by quantifying the area occupied by NTC or SYCP3-KO seeded cells. As shown in Figure 4, the area occupied by SYCP3-KO attached cells was higher at almost all the time points analyzed (15, 60 and 90 minutes), suggesting that *SYCP3* depletion enhances spreading, leading to a more stable adhesion, which might impair migration and invasion. Together, these findings indicate that *SYCP3* depletion reduces the migratory and invasive capacity of PCa cells and increases their adhesion and spreading.

### SYCP3 regulates cell morphology and F-actin cytoskeletal organization

Since our results have shown that *SYCP3* regulates invasion, adhesion and spreading we analyzed cell cytoskeleton dynamics on our cells by quantifying Actin cytoskeleton (stained with phalloidin) and nuclear areas (stained with DAPI) on cells seeded on gelatin-coated plates. Figures 4G and 4H show that SYCP3-KO cells had a larger size compared to NTC cells. Interestingly, the nuclear area was also increased in *SYCP3* depleted cells, suggesting that *SYCP3* depletion increases cell size (Figure 4I and Supplementary Figure 6). To ensure that cellular confluency did not affect nuclear and cytoskeleton area measurements we analyzed both areas by seeding at different cell densities and similar results were observed (Supplementary Figure 6).

### SYCP3 sensitizes DU145 cells to irradiation

SYCP3 activity has been previously associated with DNA damage^28^, so we decided to investigate if *SYCP3* may influence radiotherapy (RT) sensitivity on PCa cells. To analyze the effect of RT we performed survival and DNA damage analyses on SYCP3-KO and NTC DU145 cells after irradiation with photons from a Cs-137 source (Fig. 5A). As expected, we observed a decrease in the number and size of clones formed by both SYCP3-KO and NTC cells as the irradiated dose increased. However, *SYCP3* depleted cells formed more clones than NTC cells after irradiation at all tested doses (Fig. 5B-C). The experimental results have been fitted to the Linear-Quadratic model (LQ model), and the 1-sigma uncertainty of the fitting was calculated with a bootstrapping method with a sampling of N=1000. Thus, the values of the parameters of the LQ model were α_NTC_ = (0.18 ± 0.08) Gy^-1^ and β_NTC_ = (0.03 ± 0.01) Gy^-2^ for NTC and α_KO_ = (0.01 ± 0.01) Gy^-1^ and β_KO_ = (0.020 ± 0.003) Gy^-2^ for *SYCP3* depleted cells (Fig. 5D). These results suggest that *SYCP3* depletion confers resistance to RT.

**Figure 5.**
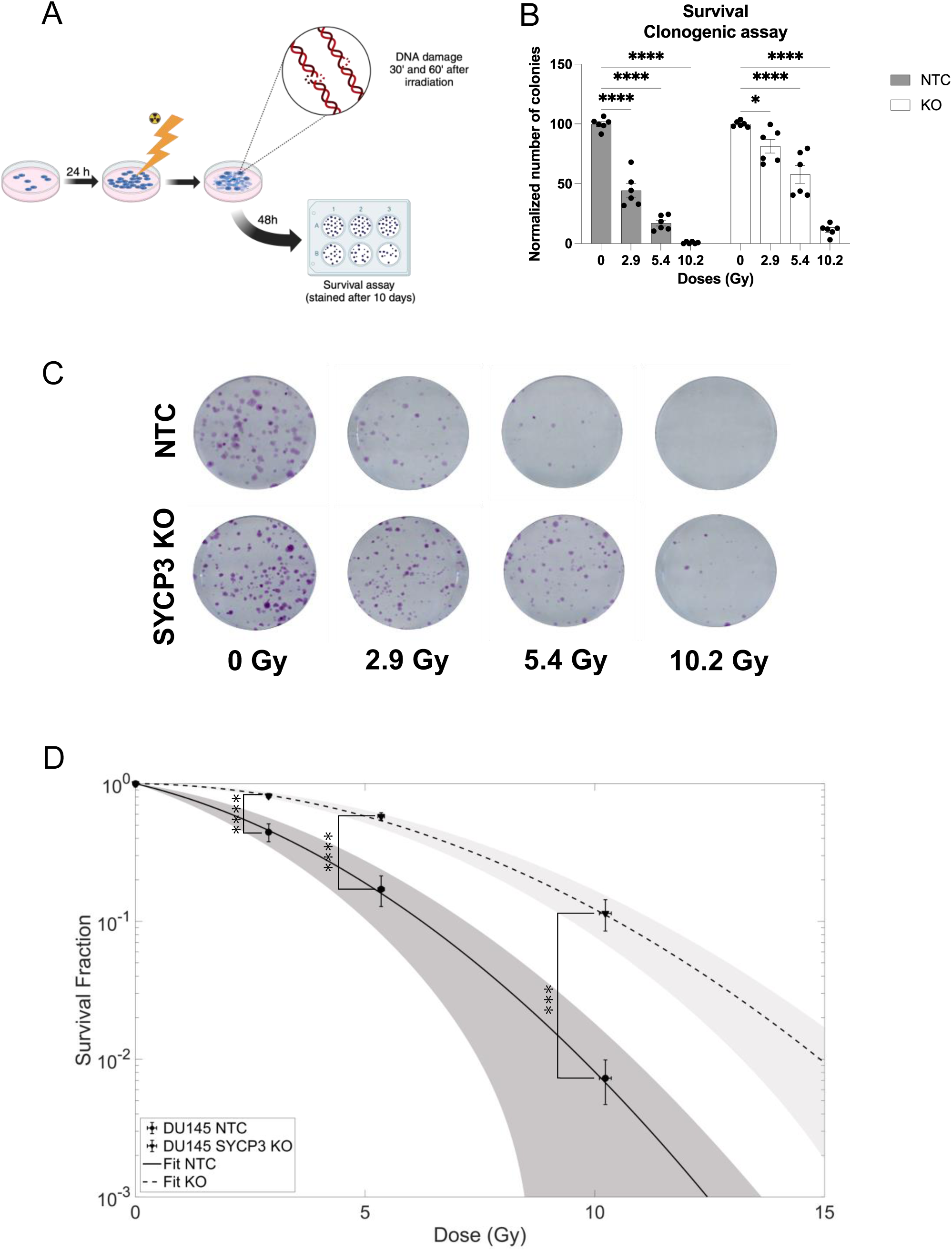
*SYCP3* depletion increases survival of PCa cells in response to RT. **A** A graphical scheme describes the irradiation plan followed in this work; cells were seeded 24 hours before irradiation. 30 and 60 minutes after irradiation the DNA damage done by the X-rays was quantified by immunofluorescence staining. 48 hours after irradiation the irradiated cells were trypsinized and reseeded at a lower seeding density for a survival assay, they were then left undisturbed in culture conditions for 10 days and, after that period, they were fixed and stained with a crystal violet solution with ethanol. **B** The stained colonies that have more than 50 cells were counted with a self-developed Matlab script and normalized to its control for each cell line (DU145 NTC and DU145 SYCP3- KO) (n=2 with triplicates).**C** Representative stained wells with the crystal violet solution from each irradiation condition and cell line. **D** The experimental results from the survival assay are shown normalized to the control and fitted to the Linear-Quadratic model (LQ model). The error bars represent the uncertainties for the survival fractions and doses, the dashed and continuous line correspond to the fitting to the LQ model and the gray areas correspond to the 1-sigma uncertainty of the fitting, this uncertainty was calculated with a bootstrapping method with a sampling of N = 1000. (n=2 with triplicates). Statistic tests: two-way ANOVA (**B, D**). *, *P* < 0.05; **, *P* < 0.01; ***, *P* < 0.001; ****, *P* < 0.0001.

DNA damage can be directly studied through the detection of protein complexes recruited to the DNA break (P-H2Ax) or involved in DNA repair processes (P53BP1). P53BP1 is known for its role facilitating non-homologous end-joining (NHEJ) and regulating homologous recombination ^29^. The number of foci generated by these proteins depends on the number of single-, double-strand breaks or the clustering of breaks. To evaluate the effect of radiation on DNA damage as well as the cellś ability to repair DNA breaks, we analyzed the density (number of foci by total DAPI area) of P-H2Ax and P53BP1 foci in SYCP3-KO and NTC cells after irradiation (Fig. 6). The cells displayed a similar P-H2Ax foci density under basal conditions. As expected, the number of P-H2Ax foci quickly increased similarly in a dose- and time-dependent manner in both cell lines after 30 minutes becoming significatively higher in SYCP3-KO cells at 10Gy (Fig. 6A and 6B). On the other hand, the density of P53BP1 foci was higher in SYCP3-KO compared to NTC cells at high doses of RT after 60 minutes (Fig. 6A, 6C and 6D), suggesting that the absence of *SYCP3* promotes the accumulation of DNA damage repair factors such as P53-BP1 following radiation treatment.

**Figure 6.**
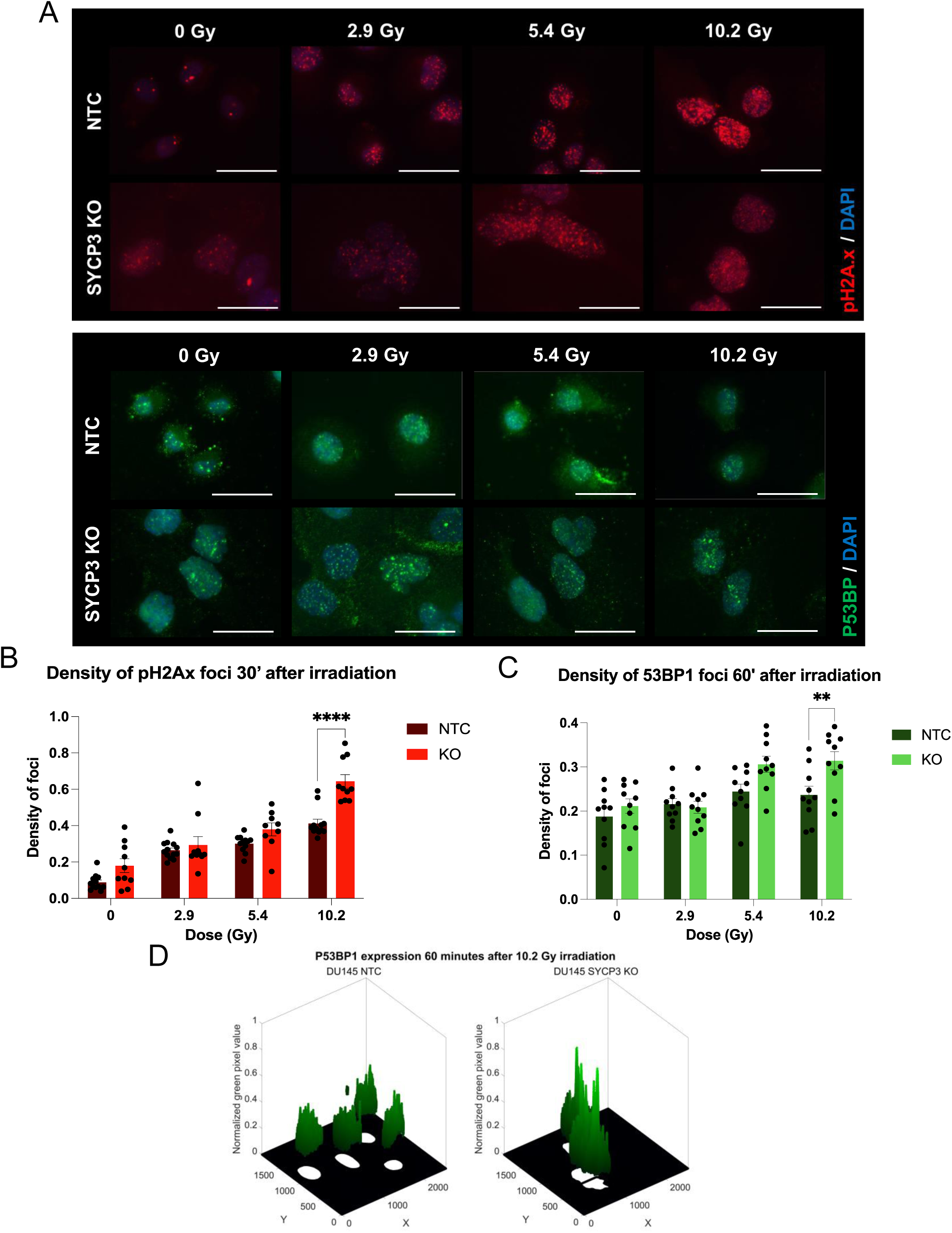
*SYCP3* depletion promotes P53BP1 related DNA repairment processes in response to RT. **A** Representative images from the DNA damage immunofluorescence for the DU145 NTC and SYCP3-KO cell lines and for each irradiating condition 30 minutes (30’) after irradiation (top, pH2Ax/DAPI) or 60 minutes (60’) after irradiation (bottom, P53BP/DAPI). In the images the two channels are merged, with the P-H2Ax staining in red, the P53BP1 in green and DAPI in blue. Scale bar: 200 µm. **B** Histogram showing the density of foci ± S.E.M. of P-H2Ax 30 minutes (min) after irradiation for the NTC (dark red) and the SYCP3-KO (light red). The density was calculated in each field by dividing the total number of foci by the total area of the nuclei (measured from the DAPI staining). The number of foci, the number of nuclei and their area were quantified with Matlab scripts. The black points correspond to the foci density for each field analyzed for each irradiation condition. **C** Histogram showing the density of foci ± S.E.M. of P53BP1 60 min after irradiation for the NTC (dark green) and the SYCP3-KO (light green). The density was calculated in each field by dividing the total number of foci by the total area of the nuclei (measured from the DAPI staining). The number of foci, the number of nuclei and their area were quantified with Matlab scripts. The black points correspond to the foci density for each field analyzed for each irradiation condition. **D** Surface plot of P53BP1 60 minutes after the 10.2 Gy irradiation for the NTC cell line (left) and the SYCP3-KO cell line (right). The *X* and *Y* axis correspond to the dimensions of the image and the *Z* axis corresponds to the normalized green pixel value, to represent the difference in the DNA damage response between both cell lines (n=2 with 10 pictures total). Statistic tests: two-way ANOVA (**B, C**). *, *P* < 0.05; **, *P* < 0.01; ***, *P* < 0.001; ****, *P* < 0.0001.

## DISCUSSION

Metastasis in PCa is a complex phenomenon involving many biochemical processes that are largely unknown ^30^. Although there are several effective therapies at the initial stages of PCa disease, metastatic castration resistant prostate cancer (mCRPC) has no cure yet^3,5^. The high incidence of this neoplasia highlights the need for the identification of new targets to be used as prognostic biomarkers and/or therapeutic targets ^1^. Our report presents a reanalysis performed on a large-scale screening specifically designed for uncovering DU145 essential invasion genes using CRISPR/Cas9 based technology^16^. The bioinformatic analysis carried out on the data previously published^16^ allowed us to identify and validate a novel gene essential for PCa cell invasive abilities. Our *in vitro* characterization of *SYCP3* unveils this gene as a regulator of mPCa cell tumor progression, being able to control invasive and migratory abilities of mPCa cells as well as PCa cell survival in response to RT.

Although *SYCP3* was first considered to be a meiosis-testis specific protein, it has been reported to be aberrantly expressed in several cancers such as human leukemia and primary cervical cancers ^24^, and tumor cell lines like hepatocellular carcinoma cell line HepG2 and the prostate cancer cell line DU145 ^31^. Its expression can be induced in culture cells by treatment with the demethylating agent 5-azacytidine indicating that physiological *SYCP3* expression is regulated by a methylation-dependent process ^28,32^ and deregulation of that process may explain its ectopic expression in cancer. Although the meiotic role of *SYCP3* is extremely well characterized ^33^, its mitotic role or its function in cancer is entirely unknown. Due to its involvement in DNA repair its expression in cancer provides a new underlying mechanism for genomic and/or chromosomal instability ^24,28^. In addition, and according to our results, *SYCP3* controls mCRPC cell dissemination and adhesive properties.

We observed differential adhesion abilities of *SYCP3* KO cells, so we focused our efforts in studying the role of SYCP3 in mPCa cell adhesion, which had not been described before. Cell adhesion to other cells and substrates is a widely known mechanism that regulates cell migration and invasion ability ^9^. Cancer cells often lose their typical adhesion patterns, becoming more invasive, which is often accompanied by smaller and more flexible shapes to enable metastasis ^20,30^. Our results showed that depletion of *SYCP3* correlates with a bigger cellular size (Fig. 4). Larger cells typically have a greater surface area providing more contact points for adhesion molecules to bind to neighboring cells and/or to the extracellular matrix (ECM) ^34^. In contrast, rounded or less spread-out cells, which tend to be smaller in their surface area, often have weaker adhesion. In addition to the larger size, *SYCP3* depleted cells displayed a higher cellular spreading, a factor often associated with larger size and stronger adhesion ^20^. Cellular spreading involves a higher number and more stable focal adhesions, which are mediated by integrins. Integrins mediate cell binding to the ECM promoting signaling through focal adhesion complexes, which anchor the cell and contribute to its size.

As shown in Figure 4F, *SYCP3* depleted cells displayed a more organized Actin cytoskeleton. This differential cytoskeleton organization can explain the higher strengthened adhesion showed by SYCP3-KO cells while less adhesive and more motile NTC cells exhibit a more dynamic Actin cytoskeleton allowing them to invade. According to our studies *SYCP3* depletion correlates with increased cell adhesion and decreased migration probably by motile structures such as lamellipodia formation, through the regulation of actin cytoskeleton. However, additional experiments need to be performed to understand how *SYCP3* controls cellular adhesion.

As mentioned, *SYCP3* is a member of the synaptonemal complex^21^. In meiosis, DSBs are deliberately induced to promote homologous recombination and ensure proper chromosome segregation. During this process *SYCP3* ensures that DSB repair occurs correctly by aligning homologous chromosomes, which serve as templates for accurate repair ^35^. Recent research has uncovered roles for *SYCP3* beyond meiosis, particularly in DNA damage and DNA repair mechanisms ^17^. Interestingly, previous reports have described its ectopic expression in tumor cells suggesting that its expression may interfere with DNA repair pathways causing improper resolution of DNA damage and accumulation of mutations^24,28^. In agreement with this, we have found that *SYCP3* expression sensitizes cells to RT.

To understand the role of *SYPC3* in the cellular response to RT we have evaluated the activity of DNA repair genes in our irradiated cellular model such as P53BP1, a key mediator of the DNA damage response, involved in recognizing and repairing DNA double-strand breaks (DSBs). Previous works have shown that P53BP1 localizes within DSBs to regulate DNA repair by facilitating non-homologous end-joining (NHEJ) and regulating homologous recombination (HR) ^29^. *SYCP3* depletion led to a higher DNA damage response dependent on P53BP1 as we see after irradiation (Fig. 6D), suggesting that irradiated *SYCP3* depleted cells promote the use of P53BP1 to increase the error prone NHEJ process facilitating DNA repair and cellular survival, while probably promoting the accumulation of mutations, inducing genomic instability. In addition, previous reports have identified a new role for *SYCP3* interfering with *BRCA2* and inhibiting the mediator role of *BRCA2* in homologous recombination (HR)^28^ . Due to this activity, *SYCP3* reduces HR efficiency and induces chromosomal instability by impairing the intrinsic repair pathway ^24^. Importantly, this inhibition of HR might be common across tumors where *SYCP3* is expressed. Our findings, in agreement with previous reports, indicate that the absence of *SYCP3*, or a compromise on its functions, may lead to a DNA damage response involving P53BP1 and/or BRCA1/2. Consequently, *SYCP3* upregulation will increase genomic instability and promote tumor diversity facilitating the development of more malignant cells, which contributes to tumor progression in a variety of cancers.

In agreement with this hypothesis, previous reports and publicly available databases confirm that *SYCP3* expression is often elevated in aggressive tumors (such as prostate, breast, and ovarian cancers) and is associated with poor outcomes and metastasis ^28^. Interestingly, *SYCP3* can promote chromosomal instability even in cancers without *BRCA2* mutations, providing a new underlying mechanism for chromosomal instability in tumor cells, probably through the activation of other DNA repair pathways regulated by P53BP1.

Therefore, our results point to *SYCP3* as a biomarker of poor prognosis and RT sensitivity for mPCa.

## CONCLUSIONS

In summary, we showed that depletion of *SYPC3* significantly reduces invasion *in vitro.* Furthermore, we have found that its overexpression significantly correlates with tumor aggressiveness and a reduced disease-free survival of PCa patients. Last, our *in vitro* analysis points out that mechanistically *SYCP3* promotes cell migration and invasion through regulation of the adhesion capacity of cells. Moreover, *SYCP3* regulates the radiosensitivity of PCa tumor cells. Taking all these results together, we highlight *SYCP3* as a promising biomarker candidate of poor prognosis for PCa metastasis and to select patient responders to RT.

## MATERIALS AND METHODS

### Cell culture and reagents

DU145 (HTB-81) cell line was obtained from ATCC and cultured in RPMI 10% FBS medium. HEK293T (CRL-3216) cells obtained from ATCC, were cultured in 10% FBS DMEM medium and were used for lentiviral production. Cell lines were authenticated by STR and routinely tested for mycoplasma using specific PCR analyses.

### Gene ontology (GO)

Genes candidates for studies using Gene ontology (GO) enrichment analyses of Figure 1, 2A-B were performed using 884 genes that significantly (pValue<0.05 and gRNA>1) reduced the invasive capacities of DU145 cells according to the MAGECK analysis previously performed ^16^ (Supplementary Table 1) and https://geneontology.org/ ^36,37^ . The most enriched biological pathways were represented using R statistical environment 4.4.2 (https://www.r-project.org/). GSEA analyses from Figure 2C-D were performed using GSEA software ^38^ and the hallmark gene set of the Human Molecular Signatures Database (MSigDB). The most enriched biological pathways were represented using R statistical environment 4.4.2 (https://www.r-project.org/).

### SYCP3 CRISPR/Cas9 knock-out production

Lenti-sgSYCP3 and lenti-sgNTC viral production was performed in HEK293T cells seeded in 10 cm tissue culture plates. A mixture of the 3 transfection plasmids (Packaging and enveloping plasmids psPAX2 (Addgene, #12260), pCMV-VSV-G (Addgene, #8454) together with specific sgRNA for *SYCP3* (Forward 5’-CACCGTAAACTGATCTTCCACAGA-3’ and Reverse 5’-aaacTCTGTGGAAGATCAGTTTAC-3’) or a control (sgNTC) (Forward 5’-CACCGCGGCTGAGGCACCTGGTTTA-3’ and Reverse 5’-AAACTAAACCAGGTGCCTCAGCCG-3’); different from the ones included in the initial screening, were mixed in a 2:1:4 ratio and transfected to the HEK293T cells using 18µL of Lipofectamine 2000 (Invitrogen). After 72h, the medium was collected, centrifuged at 2000 rpm for 5 min, and filtered with a 0.45 µm pore syringe filter. DU145-Cas9 previously engineered in the laboratory, were infected with sgSYCP3 or sgNTC in presence of polybrene (8 µg/mL). After 48h, puromycin selection (10 µg/mL) was added for 5 days. A serial dilution was performed in 96 multi-well plates to obtain single cell clones. *SYCP3* knock-out was confirmed by ICE analysis and western-blot using anti-SYCP3 antibody.

### ICE analysis

Genomic gDNA was extracted from cells using standard protocols. Genomic regions flanking the CRISPR target codon start site was PCR amplified using a Dream Taq DNA polymerase (Thermo Fisher, Waltham, Massachusetts, USA). The sequences of the PCR primers used were Forward 5’-TGTAACCACAAATACCGGGGG-3’ and Reverse 5’-CCCCCAAATAAATGAGGAAACTGT-3’. PCR products were electrophoretically separated in 1% agarose gel, followed by gel extraction (Gel and PCR Clean-up Kit, Düren, Germany) and Sanger sequencing (Genomic and Proteomic unit, UCM). DNA sequencing chromatograms were analyzed using the Inference of CRISPR Edits (ICE) free software tool (synthego.com/products/bioinformatics/crispranalysis).

### Invasion assay

Invasion was assessed in Matrigel (30µg; 45356231, Corning) coated transwells. 7.5 x 10^4^ cells were seeded in the upper chamber in serum-free medium. 5% FBS-medium was placed in the lower chamber and used as chemoattractant. After 24h, invading cells were fixed with 4% para-formaldehyde and stained with crystal violet dissolved in ethanol (0.2%). Images were taken with an Eclipse TE300 Nikon microscope and quantified using ImageJ.

### Cell adhesion and spreading assay

2,5x10^4^ cells from the DU145 NTC and SYCP3-KO cells lines were seeded by triplicate in 24 multi-well plates and kept under normal culture conditions. After 15, 30, 60 and 90 minutes the wells were washed once with PBS and stained with crystal violet with ethanol (0.2%) solution. Images were captured using a Nikon Eclipse TE300 microscope coupled to a camera. To quantify the number of cells and their areas a self-developed Matlab script was used.

### Phalloidin staining

To analyze the nucleus and cytoskeleton sizes for the DU145 NTC and SYCP3-KO cell lines, 5x10^4^ and 10^5^ cells were seeded in 24 multi-well plates on glass coverslips and maintained in normal culture conditions for 24 hours. After that period, they were washed twice with PBS, fixed with 4% para-formaldehyde (PFA) on ice for 20 minutes, washed with PBS again, permeabilized with PBS-0.5% Triton X-100 for 5 minutes at room temperature (RT) and PBS-0.1% SDS for another 5 minutes. Then, they were incubated with blocking solution (PBS-3% BSA-1.5% normal goat serum) overnight at 4°C in a humidified dark chamber.

Next, cells were incubated with phalloidin-peptide (Merck #P1951-.1MG, 1:500) and DAPI (PanReac #A4099, 1:1000) in the blocking solution for one hour at RT. After washing twice with PBS and once with distilled water, coverslips were mounted with ProLong^TM^ Gold antifade reagent (Invitrogen). Images were captured using a Nikon Eclipse TE300 fluorescence microscope coupled to a camera. To quantify the area of the cytoskeleton and the nuclei for both cell lines a Matlab script was developed.

### Irradiation

X-ray irradiation was carried out at the Instalación Radiactiva Central (IRC) facility at Universidad Complutense de Madrid. Samples from the DU145 NTC and SYCP3-KO were irradiated in a GammaCell GC10001 irradiator.

Cells from both cell lines were seeded 24 hours before the irradiation experiment, 4x10^5^ cells per condition and per cell line were seeded in 60 mm plates with four glass coverslips. The cells were seeded with normal growth media and kept in normal culture conditions until the irradiation. The irradiated doses were 2.9, 5.4 and 10.2 Gy, a non-irradiated plated from each cell line seeded in the same conditions was also taken to the facility to use as a control in the biological assays. The dosimetry was performed by the own irradiator but corrected with the appropriate attenuation factors from the material and positioning calculated by our group in the commissioning of the irradiator, done with EBT3 radiochromic films ^39^.

### Clonogenics analysis

Two days after irradiation, the remaining seeded cells in the irradiated p60 plates from the DU145 NTC and SYCP3-KO cell lines were washed twice with PBS and trypsinized with 0.05% Trypsin-EDTA (Gibco) solution. Then, cells were counted and 300 cells per irradiating condition and per cell line were seeded by triplicated in 6 multi-well plates in normal growth media. These plates were left undisturbed in normal culture conditions for 10 days. After that time, cells were washed with PBS and fixed and stained with crystal violet with ethanol (0.2%) solution. The number of colonies with more than 50 cells in each well was counted with an automatic Matlab script after scanning the plate.

### Immunofluorescence staining

30 minutes after irradiation two glass coverslips were removed, per irradiation condition, and washed twice with PBS, fixed with 4% para-formaldehyde (PFA) on ice for 20 minutes, washed with PBS, permeabilized with PBS-0.5% Triton X-100 for 5 minutes and PBS-0.1% SDS for another 5 minutes at RT. Then, they were incubated with blocking solution (PBS-3% BSA-1.5% normal goat serum) for 30 minutes at RT. Next, cells were incubated with Anti-53BP1 (P53BP1) antibody (abcam #ab175933, 1:100) and with Anti-phospho-Histone H2A.X (P-H2Ax) antibody (Merck #05-636, 1:100) in blocking solution overnight at 4°C in a humidified dark chamber. After that, the coverslips were washed three times with PBS and incubated with secondary antibody goat anti-mouse-Alexa 555 (Invitrogen #A32727, 1:500) and secondary antibody goat anti-rabbit-Alexa 488 (Invitrogen #A32731, 1:500) and DAPI (PanReac #A4099, 1:1000) in blocking solution in a humidified dark chamber at RT. After washing twice with PBS and once with distilled water, coverslips were mounted with ProLong^TM^ Gold antifade reagent (Invitrogen). Images were captured using a LEICA DMC 4500 fluorescence microscope coupled to a camera. For density of foci quantification, the number of foci per total area cover by the nuclei (positive for the DAPI staining) in each field was determined using a self-developed Matlab script.

### Mutational, expression and survival analysis using public PCa patient data

*SYCP3* mRNA expression and information about the patient and tumor GS of PCa patients was extracted from Grasso, Taylor and Varambally publicly available cohorts using CANCERTOOL database^4,25,40,41^. TCGA PRAD patient data to study *SYCP3* expression by GS was extracted from UALCAN^27^ . Mutational status data of *SYCP3* gene in different PCa cohorts of patients was analyzed using cBioportal database. To study DFS and OS of PCa patients’ data was extracted and analyzed with GePIA and cBioPortal databases, respectively ^42^.

### Statistical analysis

The results are expressed as mean value ± SEM of 1–3 independent experiments. Statistical analyses were made by ordinary t-Student tests, U-Mann Whitney, one-way ANOVA or two-way ANOVA multiple comparisons test as indicated in each experiment (p value <0.05 was considered as significant). GraphPad Prism version 8.4.2 for MacOS X, GraphPad Software, (San Diego, California USA, www.graphpad.com) has been used for representing most of the results.

### HCSC patient cohort

Slides of FFPE embedded primary tumor PCa samples were obtained from Hospital Clínico San Carlos (HCSC) Hospital Biobank (Madrid, Spain) upon informed consent and with evaluation and approval from the corresponding ethic committee (CEIC code 21/430-E). Surgical resection of primary tumor (PT) from patients with or without *de novo* metastatic disease were selected by oncologists Dr. Javier Puente Vázquez, Dr. Natalia Vidal and analyzed Pathologist Dr. Melchor Saiz-Pardo Sanz from Clínico San Carlos Hospital (Madrid, Spain). Samples were classified in two groups according to their metastatic abilities and Gleason score. Samples with de novo metastasis and 7 or above Gleason score were included in the GS≥7 group meanwhile samples without de novo metastasis and 6 or lower Gleason score were included in the GS 6 group. RNA from samples was extracted using a commercial kit, following manufacturer’s instructions (High Pure FFPE RNA Micro Kit, Roche, 4823125001). The low quantity and quality of RNA samples with less than 0.01 µg/µL of RNA and less than 0.1 of 260/230 absorbance ratio were excluded from further analysis. Levels of *SYCP3* gene expression between the GS≥7 and GS 6 groups were assessed by RT-qPCR using specific primers listed in 2Table 3 at Genetic and Genomic facility of UCM. Gene fold changes were determined by the 2-ΔΔCt algorithm. GAPDH was used as the reference gene to normalize gene expression.

## Acknowledgments

We thank the Genetic and Genomic Facility at UCM (Madrid, Spain) for the sequencing of samples. We thank the Instalacion Radioactiva Central at School of medicine UCM.

## Data and materials availability

Results are available in the main text or the supplementary materials.

---

**Supplementary Figure 1.**
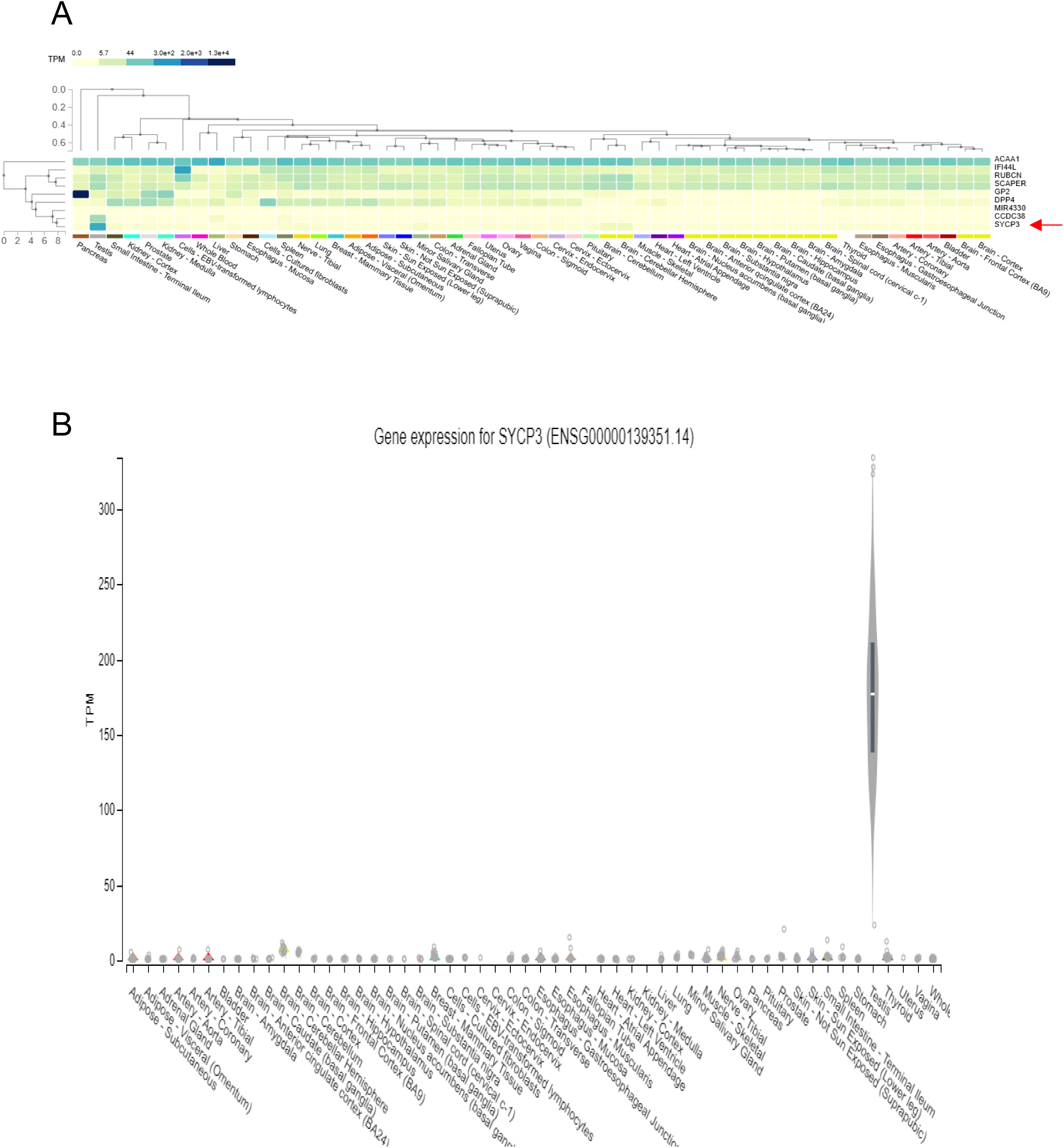
GTEx expression for the top identified genes (**A**) and SYCP3(**B**) from the genome-wide CRISPR screening in healthy tissues. The plot was generated using the GTEx Multi-Gene Query Portal (https://www.gtexportal.org/home/multiGeneQueryPage).

**Supplementary Figure 2.**
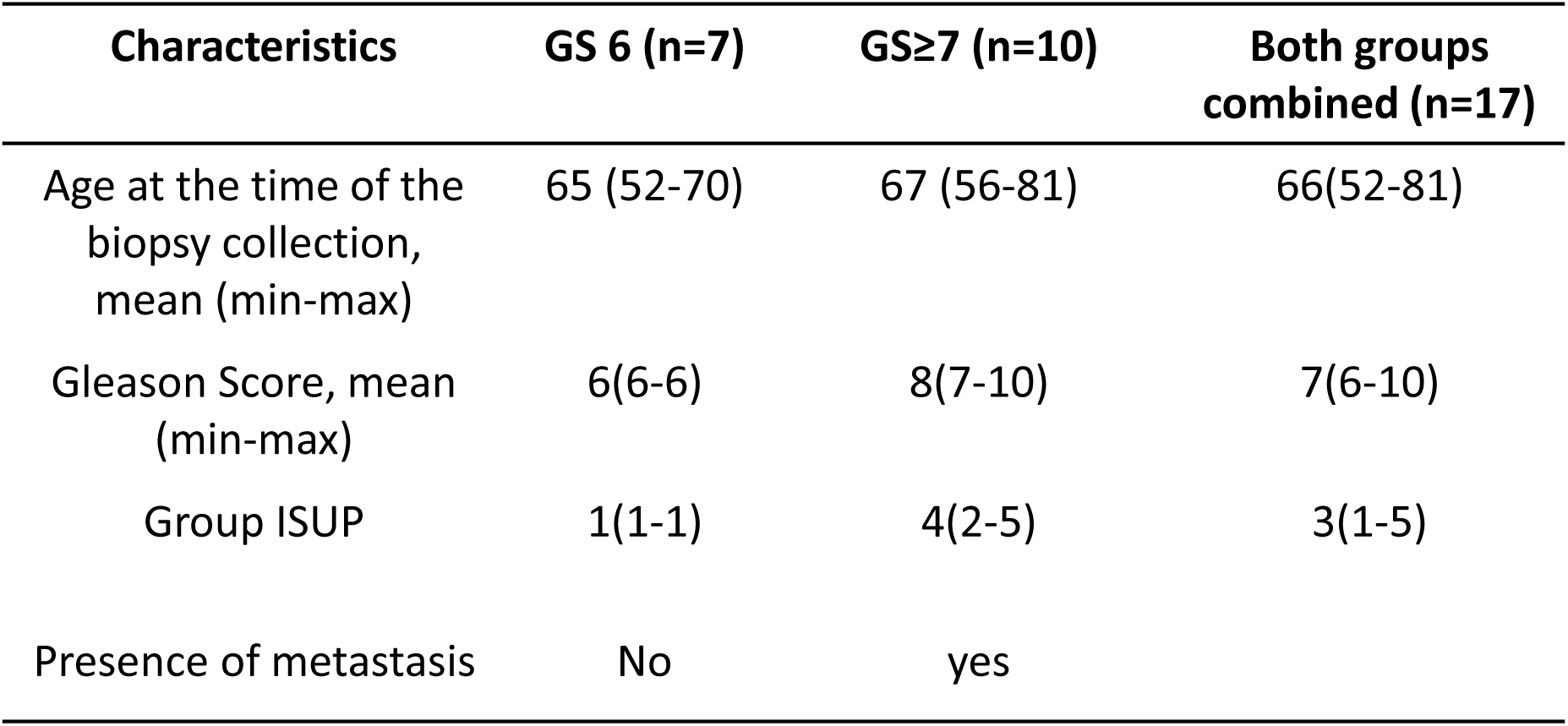
Clinical data available from the patient cohort at the San Carlos Clinical Hospital (HCSC). The characteristics of the patients and their samples are shown, divided into two groups based on their Gleason score, the presence of metastasis in the patient, and the combination of both groups (n=17 samples).

**Supplementary Figure 3.**
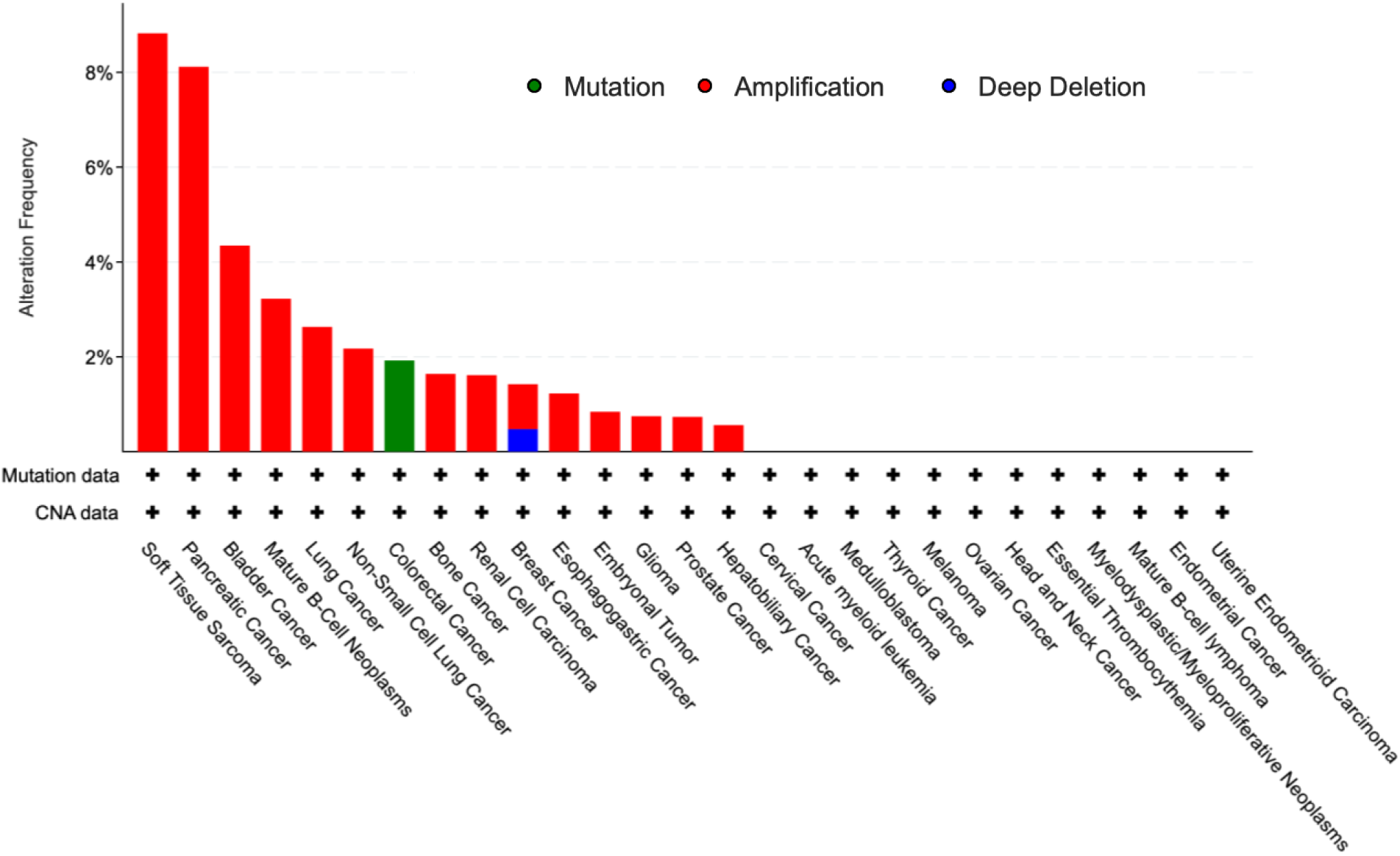
Pan-cancer analysis of *SYCP3* mutational status obtained from cBioportal.

**Supplementary Figure 4.**
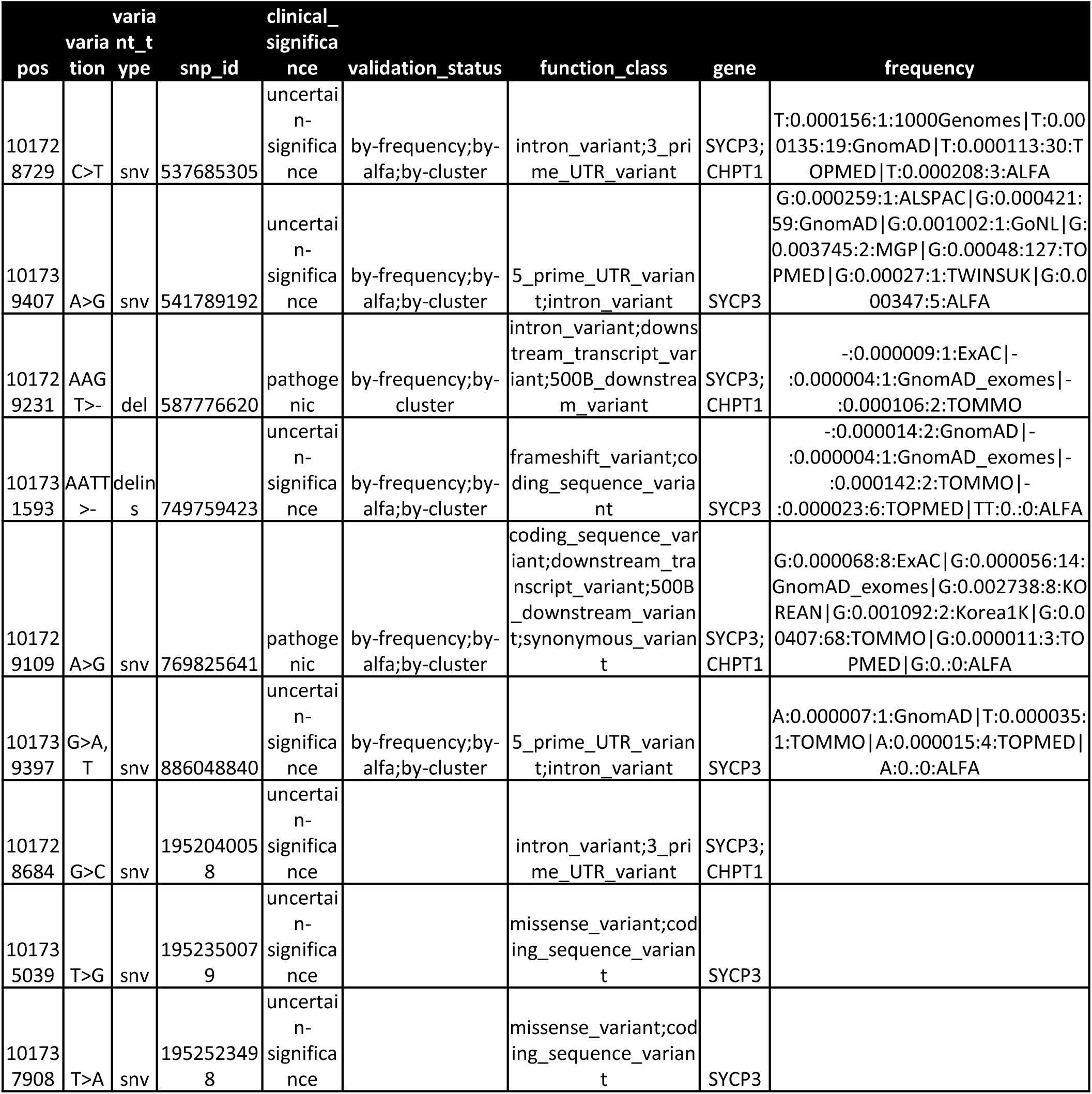
*SYCP3* genetic alterations reported in somatic cancer obtained from cBioportal.

**Supplementary Figure 5.**
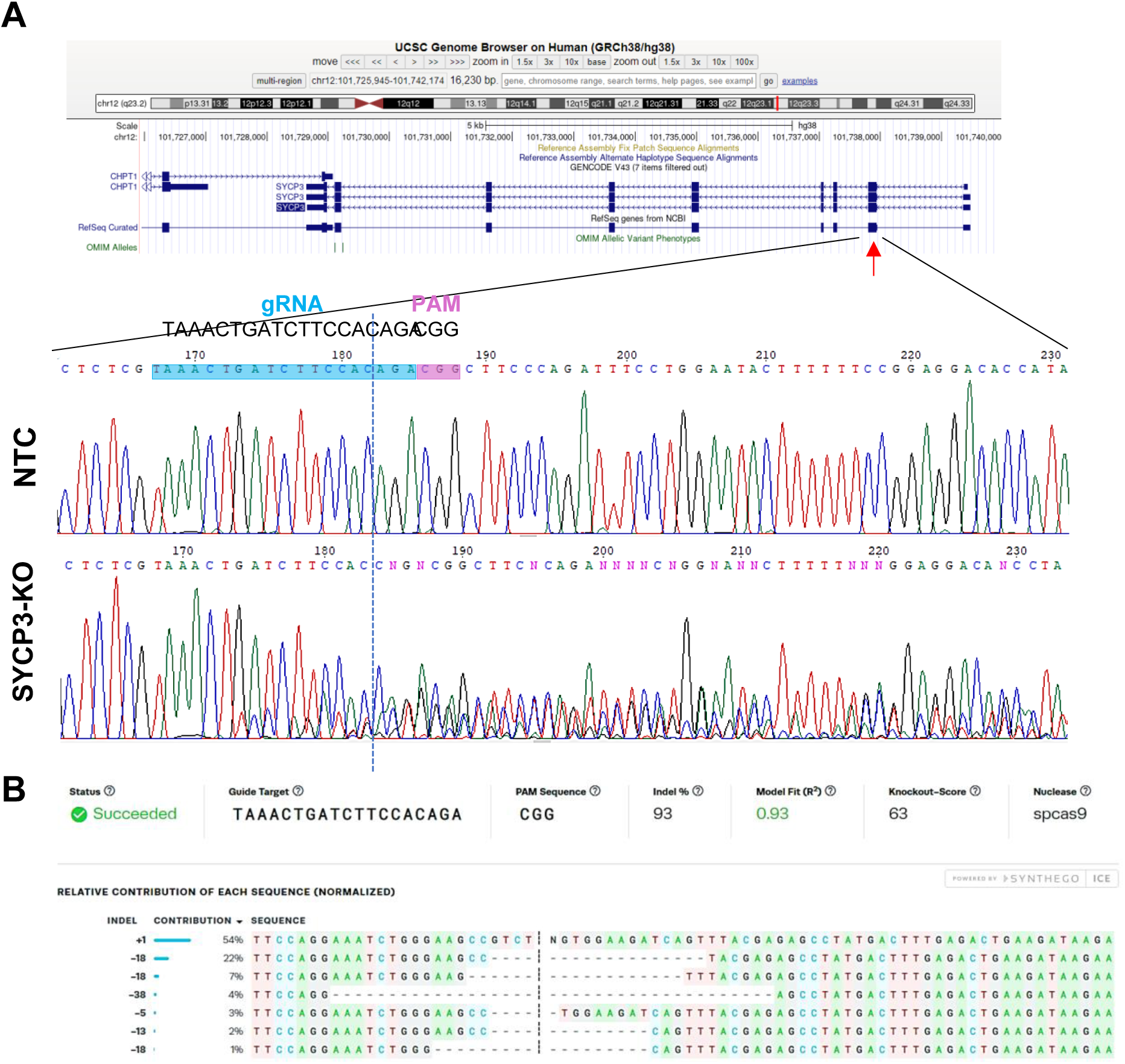
*SYCP3* genomic depletion. **A** Sanger sequencing results of NTC and SYCP3-KO cells flanking the targeted region located on the ORF exon 2 of 9. **B** The Inference of CRISPR Edits (ICE) software output of the analyses of the Sanger sequencing data on the SYCP3 gene flanking the mutated region on exon 2 of 9.

**Supplementary Figure 6.**
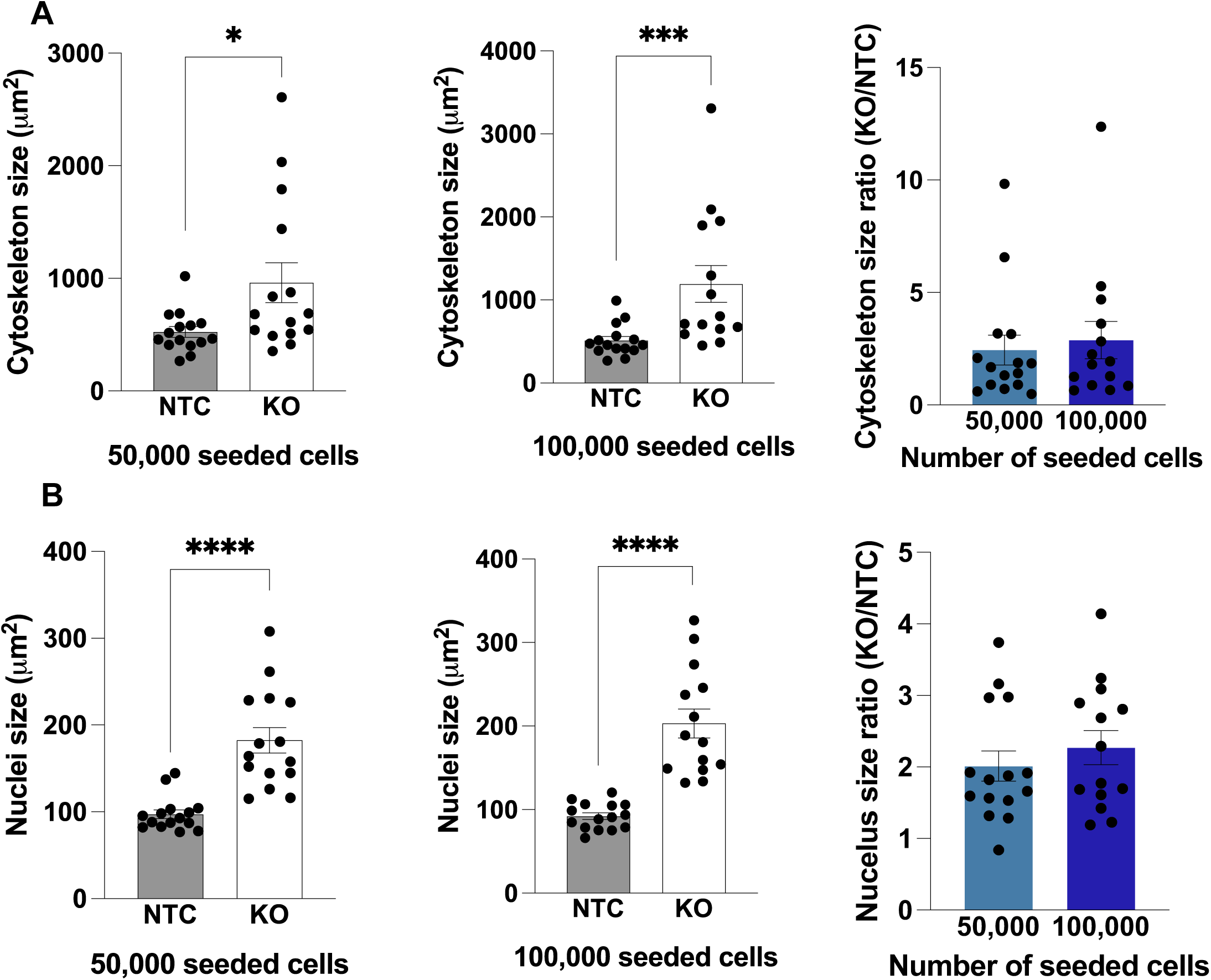
*SYCP3* depletion promotes a larger cytoskeleton and nuclei area compared to NTC cells independently of the seeded number of cells. Histograms showing the cytoskeleton **A** or nuclei area **B** value ± S.E.M. of adhered DU145 indicated cells. Statistic analysis was the Mann-Whitney test (N=2 with 15 pictures for each condition). Data were compared with the control group or as indicated, *p<0.05, ***p<0.001 and ****p<0.0001.

